# Downregulation of lysosomal trafficking in ARPE19 cells leads to decreased transfection efficiency at high passage

**DOI:** 10.1101/2023.07.26.550695

**Authors:** Erika M.S. Hood, Rachel A. Jones Lipinski, Daniel M. Lipinski

## Abstract

**PURPOSE:** ARPE19 cells are a commonly used cell culture model for the study of retinal pigment epithelial cell biology and pathologies. However, numerous studies have demonstrated that ARPE19 undergo morphologic, transcriptomic and genomic alterations over time and with increasing passage number. Herein, we explore the mechanisms underlying increased resistance to the delivery of exogenous genetic material via transfection in ARPE19 cells using mass spectrometry.

**METHODS:** ARPE19 cells (N=5 wells/reagent) were seeded in 6-well plates at passages 24 through 30. At 70% confluency an mCherry reporter construct was delivered via transfection using Lipofectamine 3000, Lipofectamine LTX, Lipofectamine Stem, or PEI (polyethylenimine) reagents. After 72 hours, transfection efficiency was quantified by fluorescence microscopy and flow cytometry. Mass spectrometry and immunofluorescence of ARPE19 cells were performed at passages 24 and 30 to evaluate altered protein synthesis and localization between passage numbers.

**RESULTS:** ARPE19 transfection showed a maximum transfection efficiency of 32.4% at P26 using Lipofectamine 3000 reagent. All lipofectamine based reagents demonstrated statistically significant decreases in transfection efficiency between passages 24 and 30. Mass spectrometry analysis revealed 18 differentially expressed proteins, including down-regulation of clathrin light chain B (CLTB) and legumain (LGMN) that was confirmed via immunofluorescence imaging, which indicated altered intracellular localization.

**CONCLUSIONS:** ARPE19 cells demonstrate passage number dependent changes in lipofectamine-based transfection efficiency. Mass spectrometry and immunofluorescence indicates the observed decrease in transfection efficiency involves the dysregulation of endocytosis and intracellular endolysosomal trafficking at later passages.

**TRANSLATIONAL RELEVANCE:** This study contributes to mounting evidence for changes in ARPE19 cell physiology with increasing passage number. This information is of value for the continued use of ARPE19 cells as a model system for RPE biology and the development of therapeutics.

## INTRODUCTION

Age related macular degeneration (**AMD**) is a leading cause of blindness in the elderly and affects an estimated 8.7% of all individuals between the ages of 45 and 85.^1^ In the earliest “dry” form of this disease, abnormal accumulations of various lipids and proteins, termed drusen, develop within Bruch’s membrane in the macula. These deposits interfere with the exchange of nutrients, waste and oxygen across Bruch’s membrane and between the retinal pigment epithelium (**RPE**) and the underlying choriocapillaris, ultimately leading to dysfunction and loss of RPE and photoreceptor cells associated with progressive loss of central vision. In approximately 10-15% of cases, disruption of nutrient and oxygen exchange following extensive drusen accumulation triggers a transition from a “dry” to neovascular or “wet” AMD, where blood vessels grow from the choroid into the retinal layers of the macula, leading to rapid vision loss due to edema and inflammation.^2–5^

ARPE19 cells are a spontaneously immortalized human RPE cell line, originally derived and characterized in 1996 and have been found to express numerous RPE cell markers such as RPE65 and CRALBP.^6^ Since then, these cells have been used to study numerous RPE cell processes, including outer segment phagocytosis, differentiation, and the effects of oxidative stress.^7–11^ ARPE19 cells have also been utilized as a cell culture model to study numerous diseases characterized by pathologies affecting RPE cells including AMD, retinitis pigmentosa, and Stargardt’s disease.^12–21^ Unfortunately, there is growing evidence to support the hypothesis that ARPE19 cells are not truly immortalized, as they demonstrate changes in morphology, abnormal karyotype, decreased viability, and signs of senescence with increasing passage number.^22–25^ This represents a major potential future problem, wherein as ARPE19 cells continue to be used, the passage number at which cells are available will inexorably increase towards the point where the cell line no longer truly recapitulates RPE cell physiology or function.

While ARPE19 cell lines have been historically useful for the study of RPE cell processes and pathologies, generally low transfection and transduction efficiencies have been a significant barrier for the use, and as such, elucidating the optimal transfection reagent for transgene delivery is an important consideration.^26,27^ In the process of comparing four commonly utilized transfection reagents over multiple passages, we identified a highly significant decrease in efficiency and explored, using a combination of immunohistochemistry and protein mass spectrometry, the mechanism underlying decreasing transfection efficiency with advancing passage number.

## MATERIALS AND METHODS

### ARPE19 cell culture

ARPE19 cells (American Type Culture Collection, Manassas, VA) were thawed at P23 from -80 °C and were initially grown in a T25cm^2^ flask in DMEM/F12 + GlutaMAX media (Gibco Life Technologies, Carlsbad, CA) containing 10% Fetal bovine serum (FBS; Gibco Life Technologies, Carlsbad, CA) and 1% Antibiotic-Antimycotic (Gibco Life Technologies, Carlsbad, CA). Upon reaching approximately 90% confluency, ARPE19 cells were passaged using TrypLE (Gibco Life Technologies, Carlsbad, CA) and seeded at a 1:5 ratio. After the initial passage, ARPE19 cells (P24-P30) were cultured in T75cm^2^ flasks with DMEM/F12 + GlutaMAX media containing 10% FBS and 1% Anti-Anti and were passaged at 95% confluency.

### Transfection

Transfection at P24-P30 was carried out in DMEM/F12 + GlutaMAX media containing 2% FBS and 1% Antibiotic-Antimycotic using Lipofectamine Stem (Thermo Fisher Scientific, Waltham, MA), Lipofectamine LTX with Plus Reagent (termed herein, Lipofectamine LTX; Thermo Fisher Scientific, Waltham, MA), Lipofectamine 3000 (Thermo Fisher Scientific, Waltham, MA) or polyethylenimine (PEI; Polysciences, PA, USA) reagents on ARPE19 cells seeded in 6-well plates at 70% confluency using 1 μg pCBA-mCherry per well and following the respective manufacturers’ protocols. Briefly, lipofectamine transfections were carried out by combining transfection reagent and DNA plasmid in OptiMEM media (Gibco Life Technologies, Carlsbad, CA), followed by an incubation period dictated by manufacturer protocols. The transfection solution is then added directly into transfection media and applied to cells. PEI transfection was carried out by combining PEI and DNA at a 2:1 ratio in transfection media, followed by a 20-minute incubation period. The PEI-DNA complex is then applied directly onto cells. Gene delivery efficiencies were quantified at 72 hours post transfection via fluorescence microscopy and flow cytometry.

### Flow Cytometry

After imaging with fluorescent microscopy, transfected and control ARPE19 cells were dissociated from the well and into a single cell suspension with TrypLE. Cells were washed once with PBS, fixed with 4% PFA for 30 minutes, and washed twice more with PBS before being resuspended in 500ul of PBS. Cell suspension was filtered with a 40um nylon mesh cell strainer (Fisher Scientific, Pittsburg, PA,) immediately before analysis with a BD LSRII flow cytometer (BD Biosciences, Franklin Lakes, NJ). 30,000 events were recorded for each well with untransfected ARPE19 cells as gating control (Supplementary Figure 1). BD FACSDiva software (BD Biosciences) was used to determine transfection efficiency.

### Histology

Untransfected control wells at passages 24 and 30 were fixed with 4% PFA for 30 minutes, permeabilized for 20 minutes with 0.2% Triton X-100 (Sigma-Aldrich, St. Louis, MO) washed four times in 0.05% Tween 20 (Sigma-Aldrich, St. Louis, MO) for 10 minutes before blocking in 0.05% Tween with 10% normal donkey serum (Sigma-Aldrich, St. Louis, MO) for one hour at room temperature. Primary antibody was applied at an appropriate concentration (Table 1) in 0.05% Tween with 2.5% normal donkey serum and incubated at 4ºC overnight. Cells were again washed four times in 0.05% Tween for 10 minutes before secondary antibody (Alexa Fluor 488, Invitrogen, Carlsbad, CA) was applied at 1:1000 dilution in 0.1% Triton X-100 and 2.5% normal donkey serum for 1 hour at room temperature. After secondary antibody removal, cells were washed three times in 0.05% Tween for 10 minutes, stained with Hoechst 33342 (1:1000 in PBS, LifeTechnologies, Eugene,OR) and washed three times with PBS for 10 minutes before being mounted. Immunofluorescence (**IF**) staining was then visualized with fluorescence confocal microscopy.

**Table 1.** Primary Antibodies Used for ARPE19 Immunofluorescence Staining.

| Antibody | Dilution | Identifier |
| --- | --- | --- |
| LGMN | 1:50 | Invitrogen #PA5-81942 |
| CLTB | 1:250 | ProteinTech #10455-1-AP |
| ZO-1 | 1:500 | Invitrogen #40-2200 |

### Mass spectrometry

#### Mass Spectrometry Sample Preparation

Untransfected ARPE19 cells were collected via trypsinization with TrypLE (Gibco Life Technologies, Carlsbad, CA) at passages 24 and 30 for mass spectrometry (**MS**). Cells were washed twice with PBS, pelleted via centrifugation at 16,100xg for 5 minutes, flash frozen, and stored at -80 °C until ready for use. Cell pellets were thawed on ice and then resuspended in 100 mM AmBic containing 20% MeCN and 2x Invitrosol (Thermo Fisher Scientific, Waltham, MA). Samples were lysed by sonication (VialTweeter; Hielscher Ultrasonics, Teltow, Germany) on ice for 25 cycles of 10 seconds on 30 seconds off. Cysteines were reduced in 5mM TCEP for 30 min at 37°C and alkylated with 10 mM iodoacetamide for 30 min at 37°C. Sequencing-grade porcine trypsin/Lys-C mix (Promega, Madison, WI) was added, and the pH was adjusted to 8.5 with NaOH (2N). Digestion proceeded for 18 h at 37 °C on a Thermomixer at 1200 rpm in the dark. Samples were acidified to below pH 3 with 10% TFA. Digested proteins were cleaned using a SolAμTM SPE plate (Thermo Fisher Scientific, Waltham, MA) on a vacuum manifold with a waste collection tray. Plate wells used for peptide clean-up were washed sequentially with LC-MS grade MeCN (1 × 200 μL) and 0.1% TFA in LC-MS grade water (1 × 200 μL). Samples were added to the wells and washed sequentially with 0.1% TFA in water (1 × 200 μL) and 0.1% FA in LC-MS grade water (1 × 100 μL). The waste collection tray was removed from the vacuum manifold and replaced with a 96-well polypropylene collection plate. Samples were eluted into the collection plate with 70% MeCN (2 × 100 μL) and the eluates were transferred to 1.5 mL microcentrifuge tubes for drying under vacuum at room temperature. Samples were resuspended by vortexing in 50 μL of LC-MS grade 2% MeCN in water. The Pierce Quantitative Fluorescent Peptide Assay (Thermo Fisher Scientific, Waltham, MA) was used to determine sample concentrations to normalize to 500 ng for each 20 µL LC-MS injection, with Thermo Scientific Pierce Peptide Retention Time Calibration Mixture added at 4 nM concentration. Equal amounts of each were combined to form a pooled QC sample.

### Mass spectrometry data analysis

Each sample was analyzed on a Thermo Scientific Orbitrap Fusion Lumos MS *via* 3 technical replicate injections using data-dependent acquisition (DDA) HCD MS2 instrument method outlined in Table S1. The technical replicates were blocked, and each block was randomized. Pooled QCs, which are a mixture of all the samples being analyzed, were analyzed at the start, end, and in between each sample block. MS data were analyzed using Proteome Discoverer 2.4 (Thermo Fisher Scientific, Waltham, MA) platform, as outlined in Table S2. Protein identifications were filtered to include only those proteins identified by two or more unique peptides identified.

Differential expression calling was performed on normalized read count data generated by Proteome Discoverer 2.4 (Thermo Fisher Scientific, Waltham, MA) using integrated Differential Expression and Pathway (iDEP) analysis (http://bioinformatics.sdstate.edu/idep/) tool.^28^ Box plots for data normalization and sample variance were assessed as part of quality control (Supplementary Figure 2). Gene enrichment analysis was subsequently performed on differentially expressed gene lists using ShinyGO 0.76 (http://bioinformatics.sdstate.edu/go/) to identify altered biological pathways between groups.

### Statistical analysis

Transfection efficiency was analyzed using one-way ANOVA with Tukey multiple comparisons test to compare transfection efficiencies between passage numbers within reagents. This was conducted with a confidence level of 95% (α = 0.05). Analysis and graphs were created using GraphPad Prism 9.

## RESULTS

### Transfection efficiency decreases with increasing passage number

To determine the relationship between increasing passage number and transfection efficiency in the ARPE19 cell line, 10,0000 cells were plated per 6-well (n=5 wells/reagent/passage) and allowed to grow to 70% confluency prior to being transfected using either Lipofectamine Stem, Lipofectamine LTX, Lipofectamine 3000, or PEI complexed with 1 μg plasmid DNA expressing a red fluorescent reporter (mCherry) transgene under the control of a ubiquitous chicken beta actin (CBA) promoter. 72 hours post-transfection cells were harvested and the transfection efficiency quantified using flow cytometry relative to untransfected controls (Supplementary Figure 1, Supplementary Figure 3).

ARPE19 transfection efficiency decreased dramatically for all transfection reagents evaluated as a function of increasing passage number, reaching statistical significance when comparing later passages (e.g. P29 or P30) to the transfection efficiency at P24 for all four reagents (Figure 1). The most effective reagent for transfecting ARPE19 cells was determined to be Lipofectamine 3000 (Figure 1A-B), with a mean transfection efficiency at P24 of 16.2% and maximum mean transfection efficiency of 32.4% occurring at P26. Transduction efficiency of ARPE19 cells with the Lipofectamine 3000 reagent decreased by P30 to 5.62%, representing a -2.88- and -5.76-fold decrease compared to P24 (p<0.0001) and P30 (p<0.0001), respectively.

**Figure 1.**
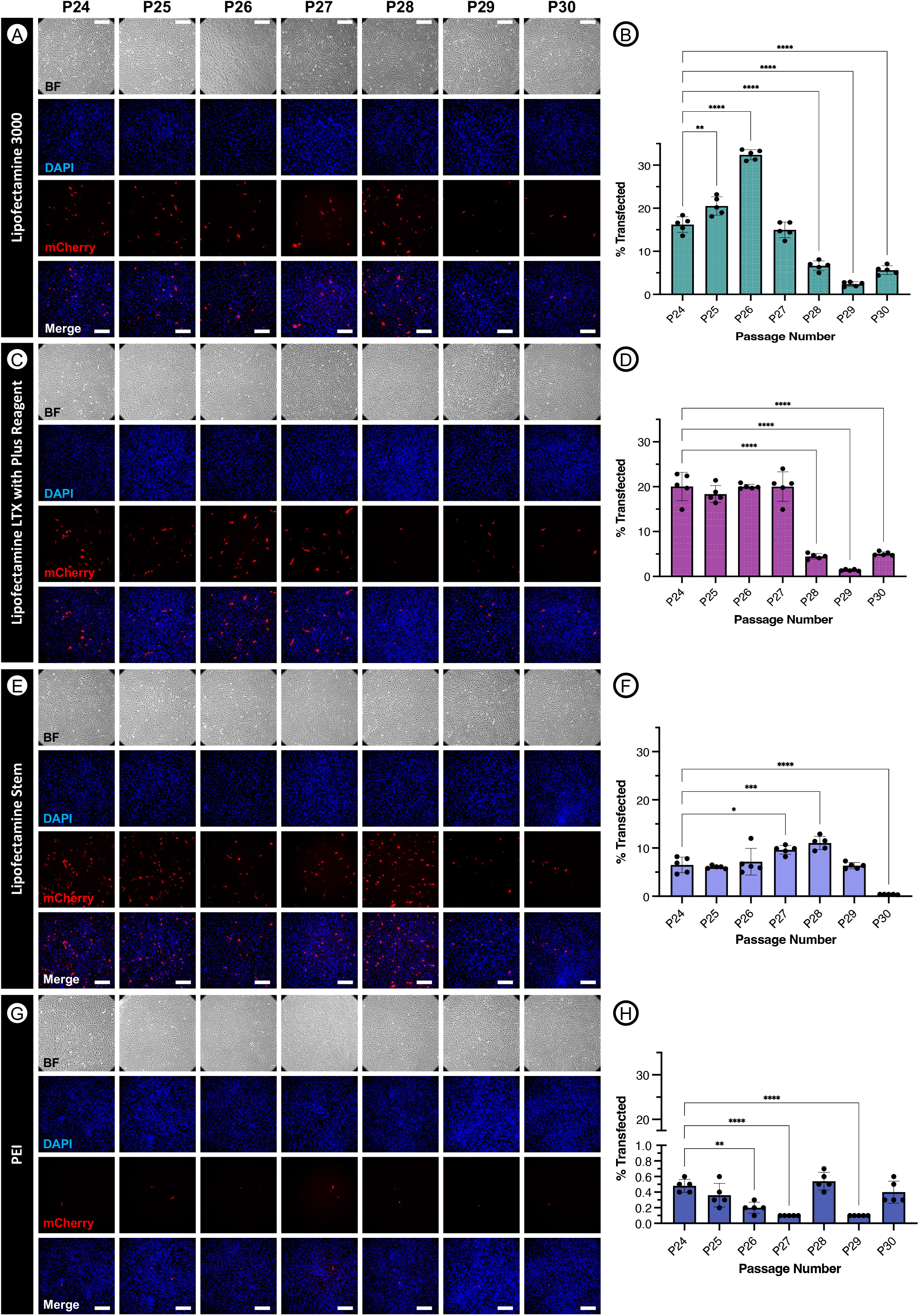
Transfection of ARPE19 cells at passages 24 through 30 using various common reagents. ARPE19 cells were transfected with 1 μg pCBA-mCherry following manufacturer protocols at 70% confluency using Lipofectamine 3000 (**A-B**), Lipofectamine LTX with Plus Reagent (**C-D**), Lipofectamine Stem (**E-F**), or PEI (**G-H**). After 72 hours, results were visualized using fluorescence microscopy and analyzed with flow cytometry. One-way ANOVA with Tukey’s multiple comparisons test p < 0.05 (*), p <0.01 (**), p<0.001 (***), p <0.0001 (****). Scale bars are approximately 200um.

Lipofectamine LTX reagent was the second most effective transfection reagent evaluated (Figure 1C-D), resulting in a maximum mean transfection efficiency of 20.04% at P24. Similarly, transfection efficiencies were significantly (p<0.0001) decreased by P30 (5.06%), representing a 3.96-fold reduction in mCherry expression observed between P24 and P30.

Transfection of ARPE19 cells with the Lipofectamine Stem reagent was only third most efficient (Figure 1E-F), reaching a mean efficiency of 6.5% at P24 and a maximum mean efficiency of 11.04% at P28, and yet demonstrated the greatest fold-change (−17.18) decrease observed between early (P24) and late-stage (P30) passages (p<0.0001).

Although transfection was observed to be least efficient with the PEI reagent (Figure 1G-H), with a maximum mean transfection efficiency of just 0.54% observed at passage 28, a statistically significant (p<0.0001; one-way ANOVA with Tukey’s multiple comparisons) -0.79-fold reduction in transfection efficiency was nevertheless observed between P24 and P29. Together, this data indicates a passage-dependent difference in transfection efficiencies, especially amongst all tested lipofectamine-based reagents.

### Mass Spectrometry of P24 versus P30 ARPE19 cells indicates down-regulation of genes involved in clathrin-mediated lysosomal transport

In order to explore the mechanistic alterations underlying the observed reduction in transfection efficiency as a result of increasing passage number, we performed MS on whole ARPE19 cell lysate harvested at P24 and P30 (N=3 biological replicates, N=9 technical replicates per time point). Quality control metrics indicated a low degree of variance between biological and technical replicates at both P24 (R_2_ = 0.956 – 0.981) and P30 (R_2_ = 0.974 – 0.979) and that transformed data was normally distributed across all samples (Supplementary Figure 2). 18 differentially expressed genes were identified as being either up-regulated (10 genes: BCAM, GNR, HEBP2, LXN, MELTF, MET, NNMT, PODXL, PRPF32A and PTGIS) or down-regulated (8 genes: ALDH16A1, CTLB, LGMN, LMCD1, MFN2, NOSTRIN, RBP1, TGM2) in P30 versus P24 ARPE19 cells (Figure 2A). Pathway analysis of differentially expressed genes revealed the upregulation of several genes involved in the positive regulation of cell-cell adhesion, epithelial cell migration and morphogenesis, and positive regulation of cell death and apoptosis, indicating that APRE19 cells tend towards greater adhesion, cell division/migration and resistance to programmed cell death with age (Figure 2B). Most interestingly, in the context of the reduced transfection efficiency observed using cationic-lipid transfection reagents (e.g. Lipofectamine stem, LTX and 3000) – which are dependent upon clathrin-mediated internalization of DNA-containing liposomes for successful transfection – was the significant (P=0.0054; -log10(FDR) = 1.11) down-regulation (−70.35 fold, P30 vs P24) of genes involved in clathrin-mediated endocytosis (e.g. clathrin light chain B; CTLB) and endo-lysosomal transport (e.g. legumain; LGMN) (Figure 2B).^29^

**Figure 2.**
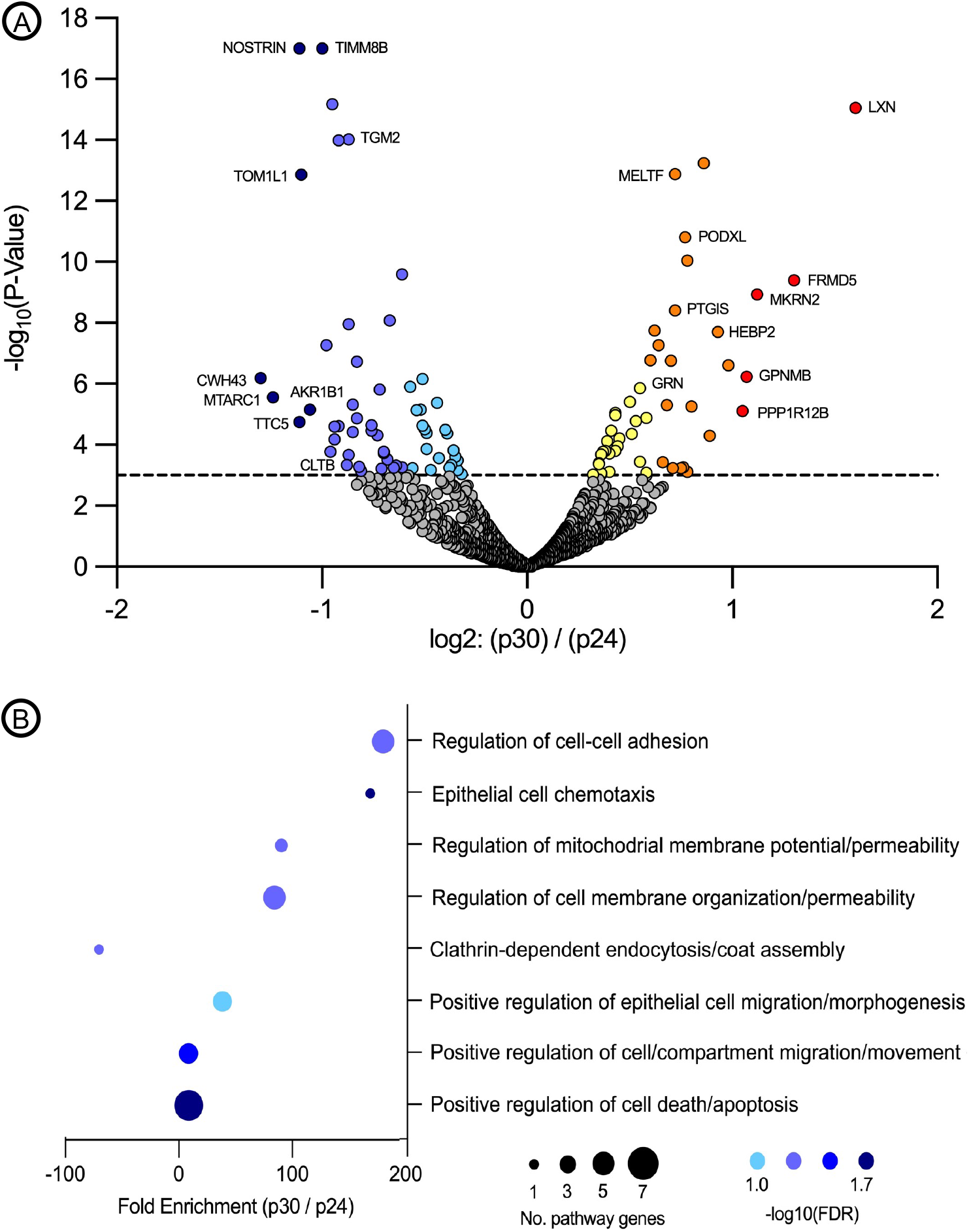
Mass spectrometry analysis of ARPE19 cells at passages 24 and 30. Volcano plot highlighting differentially expressed proteins at P30 versus P24 (**A**). Pathway analysis performed using iDEP software demonstrating pathways with major enrichment changes as a function of the number of modified genes per pathway and -log(10) false discovery rate (FDR) (**B**).

To determine whether changes in protein abundance observed by MS corresponded to observable alterations in the presence or localization of differentially regulated proteins, we performed IF staining of CLTB and LGMN in P24 and P30 ARPE19 cells and visualized the staining pattern using confocal microscopy. CLTB IF staining showed subtle changes in CLTB localization with cytoplasmic areas of lower intensity staining observable at P30 compared to P24 across all APRE19 cultures examined (Figure 3A-B, white lines). LGMN staining was similarly disrupted with a clear shift from discrete peri-nuclear localization (Figure 3C, white arrows) at P24 to a more disperse cytoplasmic staining pattern at P30 cells (Figure 3D). When combined with the previously described MS results, these data suggest dysfunction in endocytosis and lysosomal transport in ARPE19 cells at higher passage numbers. As predicted by the MS analysis, which showed no alterations in the expression levels of structural or tight junction proteins, ARPE19 cells maintained tight junctions ZO-1 at both P24 and P30, indicating decreased transfection efficiency was most likely not caused by dedifferentiation (Figure 3E-F).

**Figure 3.**
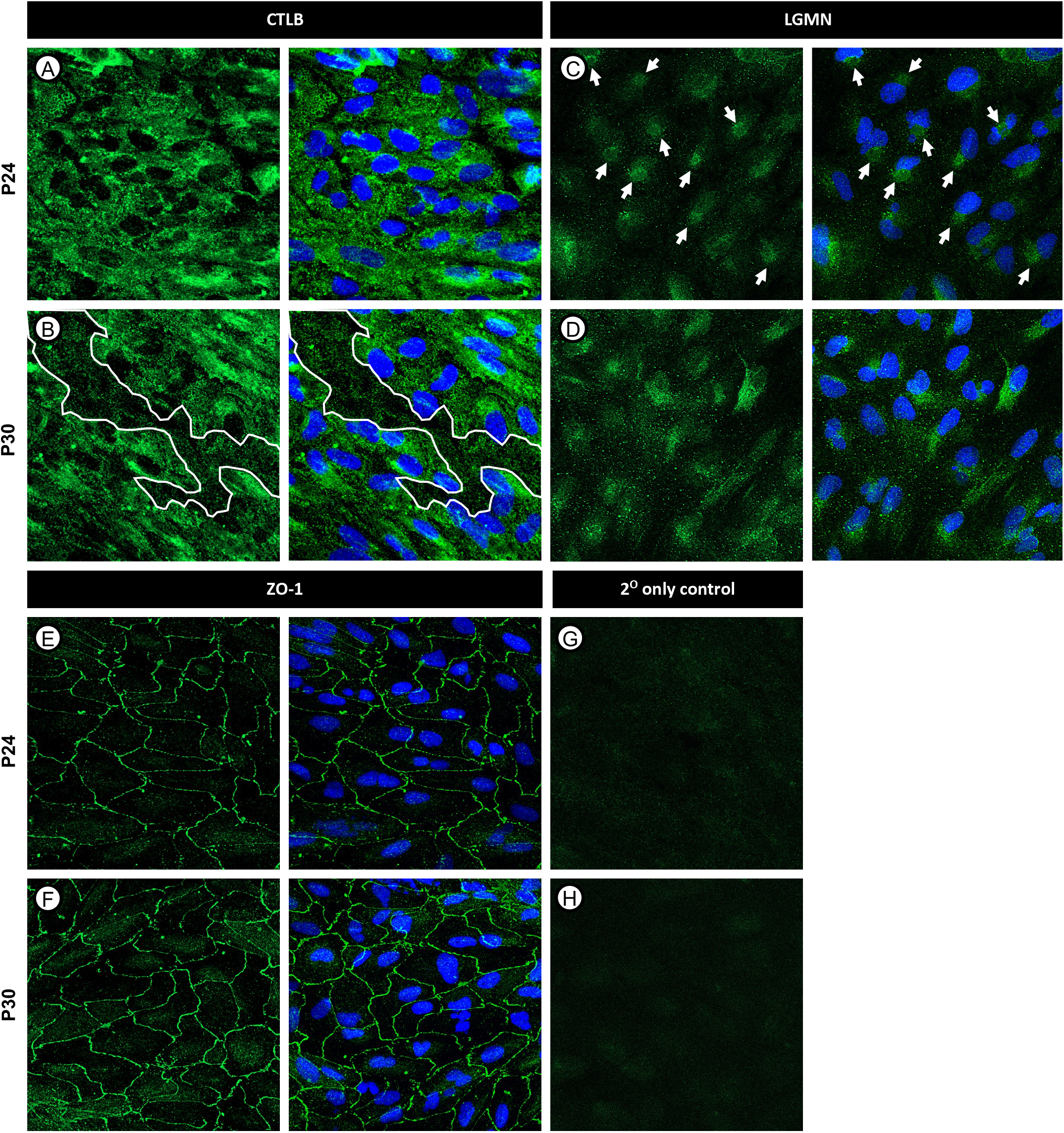
Immunofluorescence staining of CLTB, LGMN, and ZO-1 in ARPE19 cells at passages 24 and 30. ARPE19 cells were fixed and stained using immunofluorescence. CLTB staining demonstrated areas of decreased cytoplasmic staining intensity at later passages, denoted by white outlines (**A-B**). LGMN staining revealed changes in protein localization particularly surrounding the nucleus, denoted by white arrows (**C-D**). ZO-1 staining demonstrated the maintenance of tight junctions at both passages (**E-F**). Secondary antibody controls are shown in **G-H**. Scale bars are approximately 10 μm.

## DISCUSSION

ARPE19 cells have been used extensively since their isolation in 1996 and serve as an important model both for studying human RPE cell biology and the development of novel therapeutics for diseases affecting the RPE and photoreceptors, such as Leber’s congenital amaurosis type 2 (LCA2).^6^ Unlike RPE cells *in vivo*, which are highly amenable to the uptake of exogenous DNA via both non-viral (e.g. transfection) and viral (e.g. transduction) approaches, immortalized ARPE19 cells are relatively refractive to gene transfer, making them challenging to utilize.^26,27^ In this study, we set out to identify the optimal plasmid transfection reagent to efficiently delivery plasmid DNA to ARPE19 cells by screening a variety of lipid and non-lipid based transfection reagents, including polyethylenimine (PEI), Lipofectamine 3000, Lipofectamine Stem, and Lipofectamine LTX with ‘Plus’ Reagents.

As one of the primary functions of RPE cells in their native environment is the phagocytosis of shed outer segment disks, which are comprised of approximately 50% lipid by weight, we anticipated that lipofectamine-based transfection reagents that utilize lipid-DNA complexes to transverse the cell membrane would be more efficient than transfection with PEI, which relies upon endocytosis of cationic DNA-complexes across the anionic cell membrane.^30,31^ While lipofection-based transfection reagents were significantly more effective, with Lipofectamine 3000 reagent mediating maximum average transfection efficiency of 32.04 ± 1.181% versus 0.2 ± 0.071% than using PEI, the finding that transfection efficiency declines dramatically and significantly between passages 24 and 30 was unexpected. PEI transfection also demonstrated a decrease in transfection efficiency within this passage window; however, while this data may be statistically significant it is unlikely to be biologically relevant as transduction efficiencies were consistently less than 1% at all passages.

In order to explore the biological processes underlying the observed decline in transfection efficiency across all reagents with increasing passage number, we performed MS to assess relative changes in protein expression that may indicate differences in cellular physiology between ARPE19 cells at passage 24 compared to passage 30. In the context of decreased transfection efficiency, specifically with liposomal-based reagents, the most relevant observed change was a statistically significant (P=0.0054; -log10(FDR) = 1.11) down-regulation (−70.35 fold) of proteins in the clathrin-mediated endocytosis pathway, including LGMN and CLTB, at P30 compared to P24. Clathrin light chains play a critical role in clathrin-mediated endocytosis, where they form heterohexameric triskelion formations consisting of three light and three heavy clathrin chains. These triskelion formations aggregate together to form a polyhedral lattice surrounding the vesicle, and contributing towards assembly, stability and disassembly of the vesicle during uptake, and so are integral to the clathrin-mediated endocytosis process.^32–38^ As clathrin mediated-endocytosis is known to be a primary route of liposome uptake, an overall decrease in cellular levels of clathrin light chain B proteins may contribute directly to the decreased transfection efficiency at later passages, where a reduced abundance of light chains leads to decreased triskelion availability.^39^

Similarly, legumain (asparaginyl endopeptidase) is a cysteine endopeptidase commonly found within the endolysosomal pathway and is known to mediate degradation, activation, or processing of numerous substrates, playing roles in immunity and cell signaling.^40–43^ While legumain is most commonly localized within lysosomes, legumain localization outside of the endolysosomal pathway, such as in the nucleus and cytosol has been implicated in numerous pathologies such as cancer and Alzheimer’s disease.^43–48^ Additionally, legumain-deficient mice display signs of lysosomal disease and hemophagocytic syndrome.^49^ As such, unusual localization of legumain within the cell structure may be a sign cellular dysfunction.

Confocal microscopy following IF staining of APRE19 cells for CTLB and LGMN supports our MS observations, revealing areas of decreased protein expression and altered cellular localization of CLTB and LGMN at P30 compared to P24, respectively. While in this study we did not explore the mechanism through which factors associated with lysosomal trafficking are being down-regulated, that tight junction proteins such as ZO-1 – a marker of endothelial cell maturation – are not observed to be either down-regulated via MS or mis-localized on IF imaging, indicates that the underlying process does not involve the de-differentiation of the APRE19 cells as a function of increasing passage number.

In conclusion, ARPE19 cells are a commonly used model system for studying RPE biology and the development of ocular pathologies; however, their utility has come into question in recent years wherein cells exhibit changes in both morphology and viability at high passage numbers. In this study, we demonstrated that increasing passage number also affects transfection efficiency is a process involving the dysregulation of endocytosis and intracellular endolysosomal trafficking.

## ACKNOWLEDGEMENTS

The authors Michaela Pereckas and the Center for Biomedical Mass Spectrometry Research at MCW for assistance with mass spectrometry.

## FIGURE LEGENDS

**Supplementary Figure 1.**
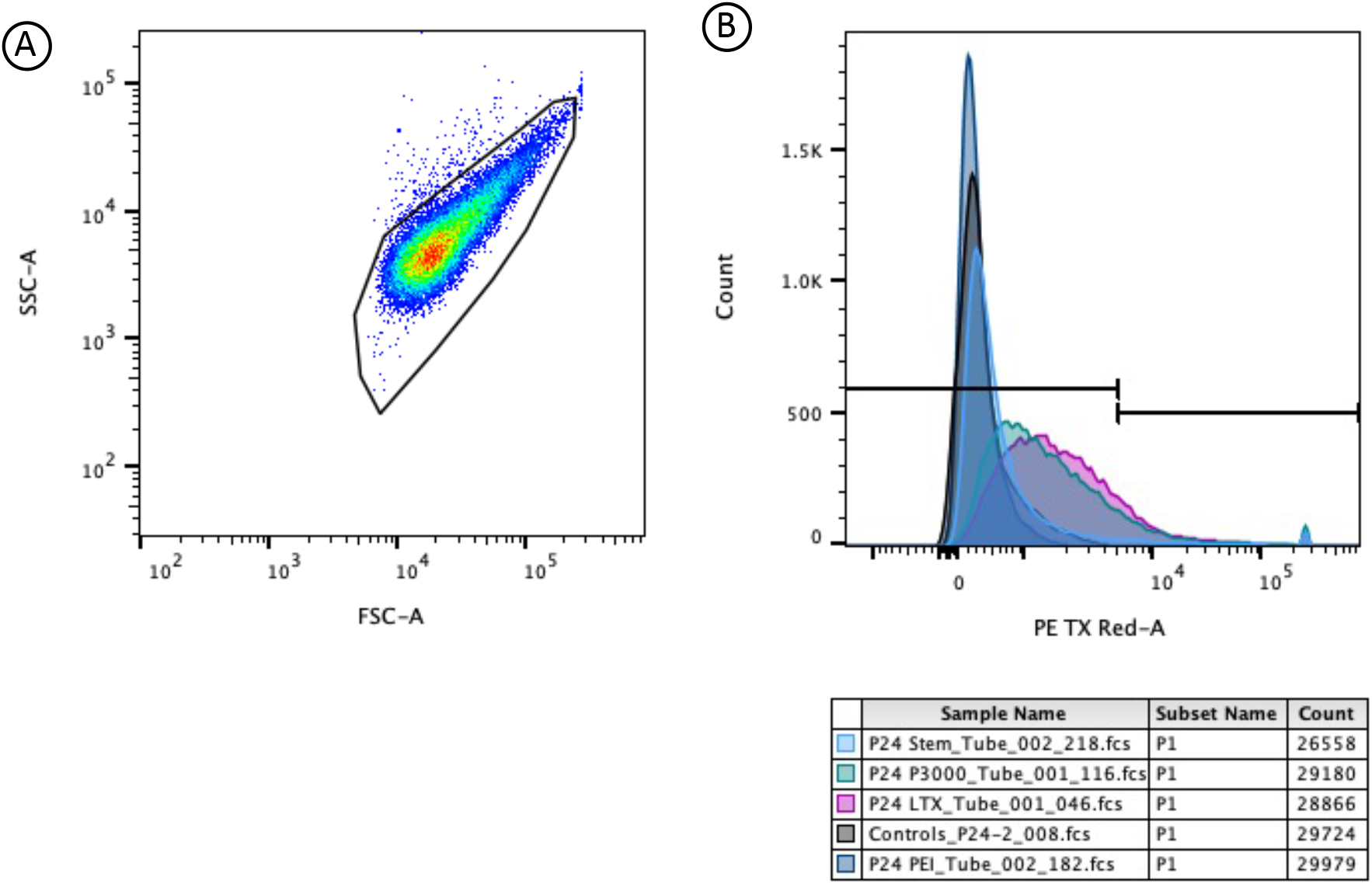
Flow cytometry gating. (**A)** Representative gating of host population for transduction efficiency evaluation. Representative transduction efficiency gating. Samples shown include n=1 for each evaluated reagent and control at passage 24.

**Supplementary Figure 2.**
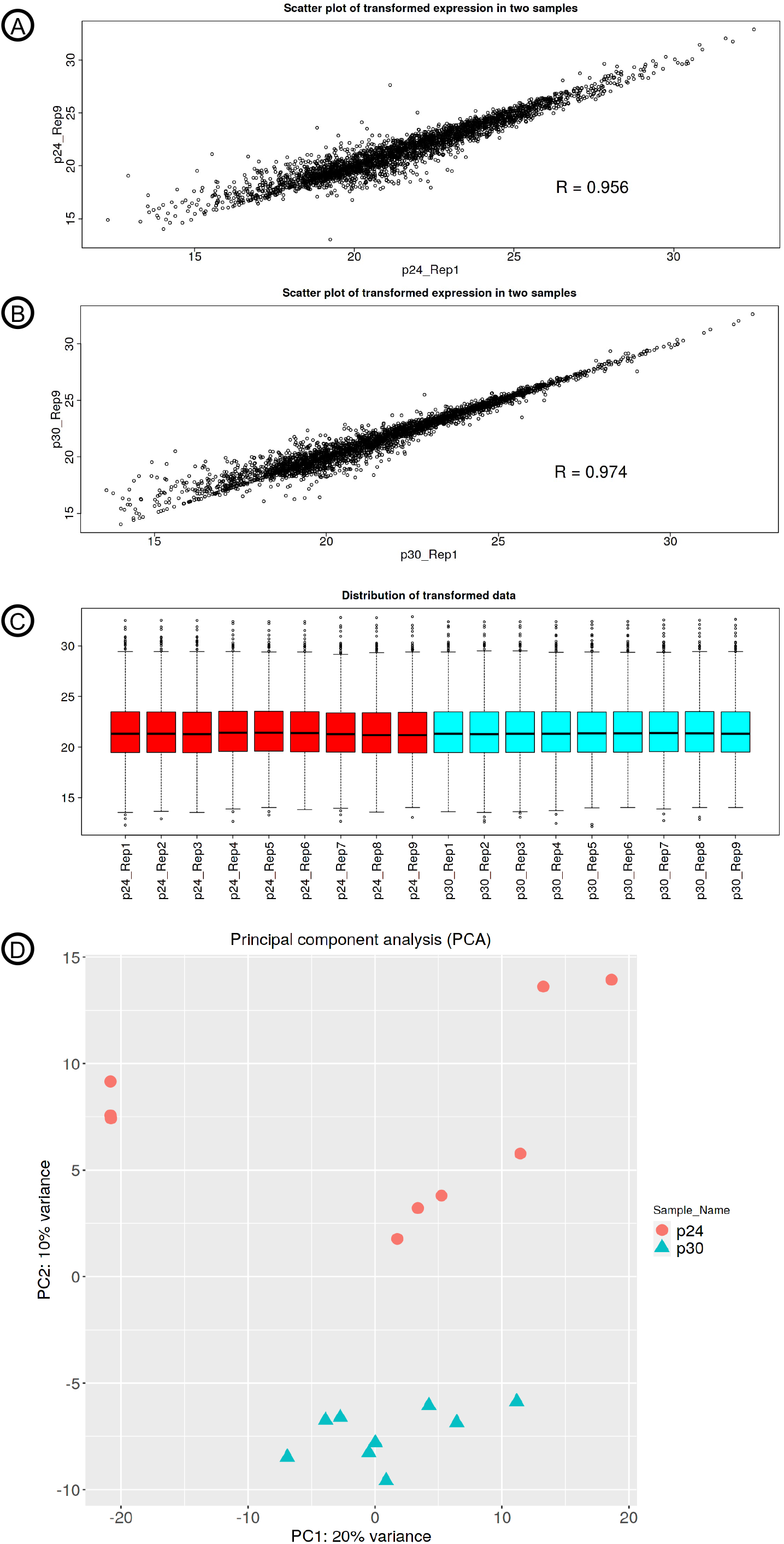
(**A)** Scatterplot visualizing log2 intensities of representative MS runs at p24 with the Pearson correlation coefficient. (**B)** Scatterplot visualizing log2 intensities of representative MS runs at P30 with the Pearson correlation coefficient. (**C)** Box and whisker plot visualizing the distribution of the normalized sample abundances. (**D)** Principal component analysis of MS data collected at P24 and P30.

**Supplementary Figure 3.**
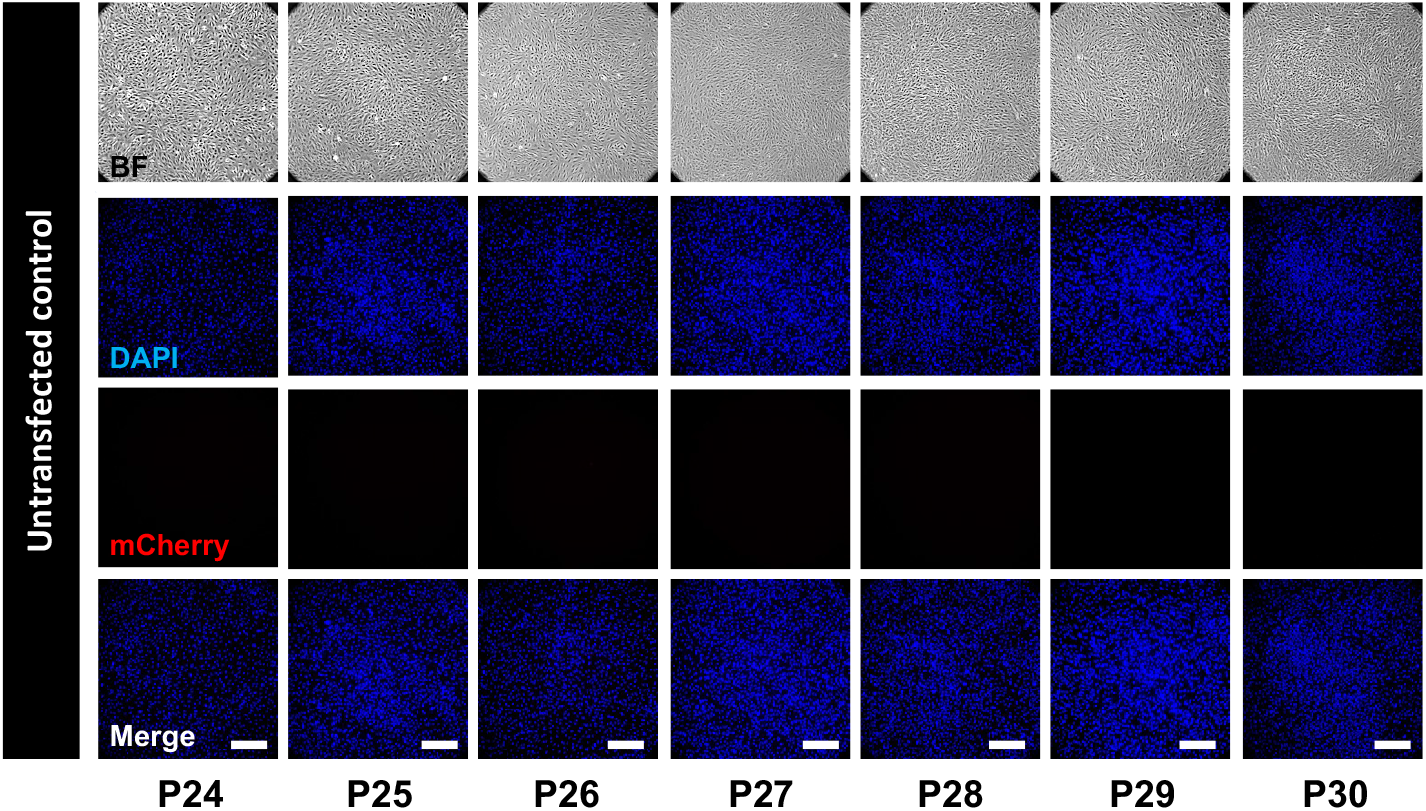
Untransfected control ARPE19 cells at passages 24 through 30. Untransfected ARPE19 cells were stained with DAPI and imaged using the same exposure and camera settings as transfected ARPE19 cells. Scale bars are approximately 200um.

## REFERENCES

1. Wong WL, Su X, Li X, et al. Global prevalence of age-related macular degeneration and disease burden projection for 2020 and 2040: A systematic review and meta-analysis. Lancet Glob Health. 2014;2(2):e106–e116. doi:10.1016/S2214-109X(13)70145-1

2. Al-Zamil WM, Yassin SA. Recent developments in age-related macular degeneration: a review. Clin Interv Aging. 2017;12:1313–1330. doi:10.2147/CIA.S143508

3. Handa JT, Bowes Rickman C, Dick AD, et al. A systems biology approach towards understanding and treating non-neovascular age-related macular degeneration. Nat Commun. 2019;10(1):3347. doi:10.1038/s41467-019-11262-1

4. Curcio CA. Soft Drusen in Age-Related Macular Degeneration: Biology and Targeting Via the Oil Spill Strategies. Invest Ophthalmol Vis Sci. 2018;59(4):AMD160-AMD181. doi:10.1167/iovs.18-24882

5. Pollreisz A, Reiter GS, Bogunovic H, et al. Topographic Distribution and Progression of Soft Drusen Volume in Age-Related Macular Degeneration Implicate Neurobiology of Fovea. Invest Ophthalmol Vis Sci. 2021;62(2):26. doi:10.1167/iovs.62.2.26

6. Dunn KC, Aotaki-Keen AE, Putkey FR, Hjelmeland LM. ARPE-19, a human retinal pigment epithelial cell line with differentiated properties. Exp Eye Res. 1996;62(2):155–169. doi:10.1006/exer.1996.0020

7. Samuel W, Jaworski C, Postnikova OA, et al. Appropriately differentiated ARPE-19 cells regain phenotype and gene expression profiles similar to those of native RPE cells. Mol Vis. 2017;23:60–89.

8. Finnemann SC, Bonilha VL, Marmorstein AD, Rodriguez-Boulan E. Phagocytosis of rod outer segments by retinal pigment epithelial cells requires alpha(v)beta5 integrin for binding but not for internalization. Proc Natl Acad Sci U S A. 1997;94(24):12932–12937. doi:10.1073/pnas.94.24.12932

9. Alizadeh M, Wada M, Gelfman CM, Handa JT, Hjelmeland LM. Downregulation of differentiation specific gene expression by oxidative stress in ARPE-19 cells. Invest Ophthalmol Vis Sci. 2001;42(11):2706–2713.

10. Kutlehria S, Sachdeva MS. Role of In Vitro Models for Development of Ophthalmic Delivery Systems. Crit Rev Ther Drug Carrier Syst. 2021;38(3):1–31. doi:10.1615/CritRevTherDrugCarrierSyst.2021035222

11. Sugano E, Edwards G, Saha S, et al. Overexpression of acid ceramidase (ASAH1) protects retinal cells (ARPE19) from oxidative stress. J Lipid Res. 2019;60(1):30–43. doi:10.1194/jlr.M082198

12. Chung EJ, Efstathiou NE, Konstantinou EK, et al. AICAR suppresses TNF-α-induced complement factor B in RPE cells. Sci Rep. 2017;7(1):17651. doi:10.1038/s41598-017-17744-w

13. Fernandez-Godino R, Bujakowska KM, Pierce EA. Changes in extracellular matrix cause RPE cells to make basal deposits and activate the alternative complement pathway. Hum Mol Genet. 2018;27(1):147–159. doi:10.1093/hmg/ddx392

14. Xu YT, Wang Y, Chen P, Xu HF. Age-related maculopathy susceptibility 2 participates in the phagocytosis functions of the retinal pigment epithelium. Int J Ophthalmol. 2012;5(2):125–132. doi:10.3980/j.issn.2222-3959.2012.02.02

15. Tseng WA, Thein T, Kinnunen K, et al. NLRP3 inflammasome activation in retinal pigment epithelial cells by lysosomal destabilization: implications for age-related macular degeneration. Invest Ophthalmol Vis Sci. 2013;54(1):110–120. doi:10.1167/iovs.12-10655

16. Yang J, Yang K, Meng X, Liu P, Fu Y, Wang Y. Silenced SNHG1 Inhibited Epithelial-Mesenchymal Transition and Inflammatory Response of ARPE-19 Cells Induced by High Glucose. J Inflamm Res. 2021;14:1563–1573. doi:10.2147/JIR.S299010

17. Bharti K, den Hollander AI, Lakkaraju A, et al. Cell culture models to study retinal pigment epithelium-related pathogenesis in age-related macular degeneration. Exp Eye Res. 2022;222:109170. doi:10.1016/j.exer.2022.109170

18. Logan S, Agbaga MP, Chan MD, et al. Deciphering mutant ELOVL4 activity in autosomaldominant Stargardt macular dystrophy. Proc Natl Acad Sci U S A. 2013;110(14):5446–5451. doi:10.1073/pnas.1217251110

19. Parmar T, Parmar VM, Perusek L, et al. Lipocalin 2 Plays an Important Role in Regulating Inflammation in Retinal Degeneration. J Immunol. 2018;200(9):3128–3141. doi:10.4049/jimmunol.1701573

20. Wang Y, Chen X, Gao X, Zhao A, Zhao C, Chen X. Variants identified by next-generation sequencing cause endoplasmic reticulum stress in Rhodopsin-associated retinitis pigmentosa. BMC Ophthalmol. 2021;21(1):371. doi:10.1186/s12886-021-02110-2

21. Li L, Jiao X, D’Atri I, et al. Mutation in the intracellular chloride channel CLCC1 associated with autosomal recessive retinitis pigmentosa. PLoS Genet. 2018;14(8):e1007504. doi:10.1371/journal.pgen.1007504

22. Pfeffer BA, Fliesler SJ. Reassessing the suitability of ARPE-19 cells as a valid model of native RPE biology. Exp Eye Res. 2022;219:109046. doi:10.1016/j.exer.2022.109046

23. Kozlowski MR. The ARPE-19 cell line: mortality status and utility in macular degeneration research. Curr Eye Res. 2015;40(5):501–509. doi:10.3109/02713683.2014.935440

24. Fasler-Kan E, Aliu N, Wunderlich K, et al. The Retinal Pigment Epithelial Cell Line (ARPE-19) Displays Mosaic Structural Chromosomal Aberrations. Methods Mol Biol. 2018;1745:305–314. doi:10.1007/978-1-4939-7680-5_17

25. Kozlowski MR. Senescent retinal pigment epithelial cells are more sensitive to vascular endothelial growth factor: implications for “wet” age-related macular degeneration. J Ocul Pharmacol Ther. 2015;31(2):87–92. doi:10.1089/jop.2014.0071

26. Gallego I, Villate-Beitia I, Martínez-Navarrete G, et al. Non-viral vectors based on cationic niosomes and minicircle DNA technology enhance gene delivery efficiency for biomedical applications in retinal disorders. Nanomedicine. 2019;17:308–318. doi:10.1016/j.nano.2018.12.018

27. Al Qtaish N, Gallego I, Villate-Beitia I, et al. Sphingolipid extracts enhance gene delivery of cationic lipid vesicles into retina and brain. Eur J Pharm Biopharm. 2021;169:103–112. doi:10.1016/j.ejpb.2021.09.011

28. Ge SX, Son EW, Yao R. iDEP: an integrated web application for differential expression and pathway analysis of RNA-Seq data. BMC Bioinformatics. 2018;19(1):534. doi:10.1186/s12859-018-2486-6

29. Rejman J, Bragonzi A, Conese M. Role of clathrin- and caveolae-mediated endocytosis in gene transfer mediated by lipo- and polyplexes. Mol Ther. 2005;12(3):468–474. doi:10.1016/j.ymthe.2005.03.038

30. Godbey WT, Barry MA, Saggau P, Wu KK, Mikos AG. Poly(ethylenimine)-mediated transfection: a new paradigm for gene delivery. J Biomed Mater Res. 2000;51(3):321–328. doi:10.1002/1097-4636(20000905)51:3<321:p:aid-jbm5>3.0.co;2-r

31. Fliesler SJ, Anderson RE. Chemistry and metabolism of lipids in the vertebrate retina. Prog Lipid Res. 1983;22(2):79–131. doi:10.1016/0163-7827(83)90004-8

32. Das J, Tiwari M, Subramanyam D. Clathrin Light Chains: Not to Be Taken so Lightly. Front Cell Dev Biol. 2021;9:774587. doi:10.3389/fcell.2021.774587

33. Silveira LA, Wong DH, Masiarz FR, Schekman R. Yeast clathrin has a distinctive light chain that is important for cell growth. J Cell Biol. 1990;111(4):1437–1449. doi:10.1083/jcb.111.4.1437

34. Huang KM, Gullberg L, Nelson KK, Stefan CJ, Blumer K, Lemmon SK. Novel functions of clathrin light chains: clathrin heavy chain trimerization is defective in light chain-deficient yeast. J Cell Sci. 1997;110 (Pt 7):899–910. doi:10.1242/jcs.110.7.899

35. Wang J, Virta VC, Riddelle-Spencer K, O’Halloran TJ. Compromise of clathrin function and membrane association by clathrin light chain deletion. Traffic. 2003;4(12):891–901. doi:10.1046/j.1600-0854.2003.00144.x

36. Ybe JA, Perez-Miller S, Niu Q, Coates DA, Drazer MW, Clegg ME. Light chain C-terminal region reinforces the stability of clathrin heavy chain trimers. Traffic. 2007;8(8):1101–1110. doi:10.1111/j.1600-0854.2007.00597.x

37. Dannhauser PN, Platen M, Böning H, Ungewickell H, Schaap IAT, Ungewickell EJ. Effect of clathrin light chains on the stiffness of clathrin lattices and membrane budding. Traffic. 2015;16(5):519–533. doi:10.1111/tra.12263

38. Ferreira F, Foley M, Cooke A, et al. Endocytosis of G protein-coupled receptors is regulated by clathrin light chain phosphorylation. Curr Biol. 2012;22(15):1361–1370. doi:10.1016/j.cub.2012.05.034

39. Cui S, Wang B, Zhao Y, et al. Transmembrane routes of cationic liposome-mediated gene delivery using human throat epidermis cancer cells. Biotechnol Lett. 2014;36(1):1–7. doi:10.1007/s10529-013-1325-0

40. Chen JM, Dando PM, Rawlings ND, et al. Cloning, isolation, and characterization of mammalian legumain, an asparaginyl endopeptidase. J Biol Chem. 1997;272(12):8090–8098. doi:10.1074/jbc.272.12.8090

41. Chen JM, Dando PM, Stevens RA, Fortunato M, Barrett AJ. Cloning and expression of mouse legumain, a lysosomal endopeptidase. Biochem J. 1998;335 (Pt 1)(Pt 1):111–117. doi:10.1042/bj3350111

42. Dall E, Brandstetter H. Structure and function of legumain in health and disease. Biochimie. 2016;122:126–150. doi:10.1016/j.biochi.2015.09.022

43. Solberg R, Lunde NN, Forbord KM, Okla M, Kassem M, Jafari A. The Mammalian Cysteine Protease Legumain in Health and Disease. Int J Mol Sci. 2022;23(24). doi:10.3390/ijms232415983

44. Haugen MH, Johansen HT, Pettersen SJ, et al. Nuclear legumain activity in colorectal cancer. PLoS One. 2013;8(1):e52980. doi:10.1371/journal.pone.0052980

45. Gawenda J, Traub F, Lück HJ, Kreipe H, von Wasielewski R. Legumain expression as a prognostic factor in breast cancer patients. Breast Cancer Res Treat. 2007;102(1):1–6. doi:10.1007/s10549-006-9311-z

46. Basurto-Islas G, Grundke-Iqbal I, Tung YC, Liu F, Iqbal K. Activation of asparaginyl endopeptidase leads to Tau hyperphosphorylation in Alzheimer disease. J Biol Chem. 2013;288(24):17495–17507. doi:10.1074/jbc.M112.446070

47. Zhang Z, Song M, Liu X, et al. Cleavage of tau by asparagine endopeptidase mediates the neurofibrillary pathology in Alzheimer’s disease. Nat Med. 2014;20(11):1254–1262. doi:10.1038/nm.3700

48. Pivtoraiko VN, Stone SL, Roth KA, Shacka JJ. Oxidative stress and autophagy in the regulation of lysosome-dependent neuron death. Antioxid Redox Signal. 2009;11(3):481–496. doi:10.1089/ars.2008.2263

49. Chan CB, Abe M, Hashimoto N, et al. Mice lacking asparaginyl endopeptidase develop disorders resembling hemophagocytic syndrome. Proc Natl Acad Sci U S A. 2009;106(2):468–473. doi:10.1073/pnas.0809824105

